# Differential Redox Regulation and Antioxidant Dynamics in Tomato Fruits under Mercury Stress

**DOI:** 10.64898/2026.05.26.727803

**Authors:** Ramzi Murshed, Sanders Junglee, Huguette Sallanon, Laurent Urban, Félicie Lauri

**Affiliations:** Université d’Avignon et des Pays de Vaucluse, Institut Agrosciences, Environnement et Santé (AgES)

**Keywords:** oxidative stress, antioxidant enzymes, ascorbate–glutathione cycle, mercury toxicity, *Solanum lycopersicum*

## Abstract

The present study investigated the impact of mercury exposure on plant water status, oxidative stress markers, and antioxidant defense systems in leaves and fruits of tomato (Solanum lycopersicum L. cv. Micro-Tom). Plants were exposed to increasing concentrations of HgCl_2_ for 24 and 48 h. Mercury treatment led to a significant reduction in predawn leaf water potential, whereas other water-related parameters in both leaves and fruits remained largely unaffected.

Oxidative stress was predominantly observed in leaves, as indicated by elevated hydrogen peroxide (H_2_O_2_) and malondialdehyde (MDA) levels, while fruit tissues showed no significant accumulation of these oxidative markers. In contrast, fruits exhibited a marked activation of antioxidant defenses, including increased activities of superoxide dismutase (SOD) and catalase (CAT), along with concentration-dependent modulation of ascorbate–glutathione cycle enzymes (APX, MDHAR, DHAR, and GR) and their corresponding transcript levels.

Alterations in the ascorbate pool, reflected by changes in reduced ascorbate (AsA) and dehydroascorbate (DHA), further indicated a dynamic regulation of cellular redox status in response to mercury exposure. Collectively, these findings demonstrate that mercury induces tissue-specific oxidative responses and rapidly activates antioxidant mechanisms in fruits, thereby contributing to the maintenance of redox homeostasis and protection against oxidative damage.

## 1. Introduction

Heavy metal contamination of soils has emerged as a critical environmental constraint with far-reaching implications for agricultural sustainability and food safety, primarily due to the potential transfer of toxic elements into trophic systems (Golia, 2023). Among these contaminants, mercury (Hg) is particularly problematic owing to its persistence, high mobility, and strong bioaccumulative behavior in both terrestrial and aquatic compartments. Agricultural practices may inadvertently introduce mercury into soils through amendments such as sewage sludge, mineral fertilizers, pesticides, and organic residues, while anthropogenic inputs, including industrial effluents and domestic discharges, further intensify Hg loading in aquatic environments (Patra and Sharma, 2000; Gworek et al., 2020; Huang et al., 2015).

At the cellular and molecular levels, mercury exerts pronounced phytotoxic and genotoxic effects (Israr et al., 2011). Its reactivity with biomolecules enables mercuric ions to interact with nucleic acids, leading to the formation of stable covalent adducts with DNA (Sharma and Talukder, 1989), and to the induction of chromosomal instability, including sister chromatid exchanges (Sánchez Alarcón et al., 2021). In addition, mercury disrupts key physiological processes in plants, impairing photosynthetic performance, water relations, and pigment biosynthesis, while simultaneously promoting membrane lipid peroxidation and metabolic dysfunction (Mei et al., 2021; Cho and Park, 2000; Chakraborty and Choudhury, 2023).

Exposure to elevated concentrations of heavy metals is commonly associated with the onset of oxidative stress, a condition arising from an imbalance between reactive oxygen species (ROS) generation and detoxification (Yamamoto et al., 1997). Under stress conditions, perturbations in chloroplast photochemistry enhance the leakage of excess excitation energy toward oxygen, resulting in the formation of ROS such as superoxide radicals, singlet oxygen, hydrogen peroxide, and hydroxyl radicals (Mittler, 2002). These highly reactive species can severely compromise cellular integrity by inducing lipid peroxidation, protein oxidation, enzyme inhibition, pigment degradation, and genomic instability (Halliwell, 2006; Mandal et al., 2022).

To maintain redox homeostasis, plants have evolved a complex and highly coordinated antioxidant network. Metal toxicity becomes critical when intracellular concentrations exceed the sequestration and detoxification capacities of plant tissues (Sinha et al., 1996). The primary defense system consists of enzymatic components, where superoxide dismutase (SOD) catalyzes the dismutation of superoxide radicals into hydrogen peroxide, which is subsequently detoxified by catalase (CAT) or ascorbate peroxidase (APX) (Fridovich, 1986; Willekens et al., 1995; Bowler et al., 1992).

A central hub of redox regulation is the ascorbate–glutathione cycle, which operates through a tightly regulated sequence of redox reactions. In this pathway, APX mediates the oxidation of ascorbate (AsA) to monodehydroascorbate (MDHA), which is either enzymatically reduced back to AsA via monodehydroascorbate reductase (MDHAR) or spontaneously converted into dehydroascorbate (DHA). DHA is subsequently reduced to AsA by dehydroascorbate reductase (DHAR) using reduced glutathione (GSH) as an electron donor, while glutathione reductase (GR) regenerates GSH from its oxidized form (GSSG) in an NADPH-dependent manner (Foyer et al., 1994; Noctor and Foyer, 1998). This cycle plays a pivotal role in sustaining cellular redox buffering capacity and preventing ROS overaccumulation. Complementary to enzymatic defenses, low-molecular-weight antioxidants, including ascorbate, glutathione, tocopherols, and carotenoids, contribute to the fine-tuning of redox balance and oxidative stress mitigation (Foyer et al., 2005).

Despite extensive recognition of heavy metal toxicity, the mechanistic basis of mercury-induced phytotoxicity remains incompletely resolved, particularly with respect to its interaction with oxidative metabolism. The scarcity of integrative data hampers a precise evaluation of the contribution of oxidative stress to mercury toxicity in plants. Therefore, the present study aims to elucidate whether oxidative stress constitutes a central component of mercury-induced damage in tomato plants, and to characterize the dynamic responses of enzymatic and non-enzymatic antioxidant systems under Hg exposure.

## 2. Materials and Methods

### 2.1. Plant material and experimental design

Tomato plants (Solanum lycopersicum L., cv. Micro-Tom) were cultivated in 4 L plastic pots containing a peat–vermiculite mixture (1:1, v/v) under controlled growth chamber conditions. Environmental parameters were maintained at 60 ± 5% relative humidity, with a day/night temperature regime of 25/20 °C and a 16/8 h photoperiod under a light intensity of 300 µmol m^−2^ s^−1^ provided by fluorescent lamps.

Mercury treatments were initiated during the fruit development stage by supplementing the growth substrate with HgCl_2_ at concentrations of 2, 5, 10, and 20 ppm. Untreated plants were used as controls. Fruits at the mature green stage (35–40 days post-anthesis) were collected after 24 and 48 h of exposure. Immediately after harvest, samples were snap-frozen in liquid nitrogen, finely ground, and stored at ^−^80 °C until further analyses.

### 2.2. Determination of water status parameters

Predawn leaf water potential (LΨw) was measured using a pressure chamber following the approach of Scholander et al. (1965). Fruit water potential (FΨw) was determined using a dew point microvoltmeter (HR-33T) equipped with C-52 chambers (Wescor Inc., USA), while fruit osmotic potential (FΨO) was assessed using a vapor pressure osmometer (VAPRO, Wescor).

To evaluate water flux, plants were excised approximately 1 cm above the substrate surface. Fruit fresh weight (FW) was recorded immediately after sampling, whereas dry weight (DW) was obtained after oven-drying at 70 °C for 72 h. Fruit water content (FWC) was calculated according to the equation:FWC = (FW ^−^ DW) / FW × 100.

### 2.3. Quantification of hydrogen peroxide

Hydrogen peroxide (H_2_O_2_) levels were quantified according to a microplate-adapted protocol. Approximately 0.25 g of frozen fruit tissue was homogenized in 1 mL of cold 0.1% (w/v) trichloroacetic acid (TCA) and centrifuged at 12,000 × g for 15 min at 4 °C.

Aliquots of the supernatant (100 µL) were mixed with potassium phosphate buffer (10 mM, pH 7.0) and potassium iodide (KI), and the reaction mixture was incubated prior to absorbance measurement at 390 nm using a microplate reader. Hydrogen peroxide concentration was calculated based on a standard calibration curve generated using known H_2_O_2_ concentrations.

### 2.4. Lipid peroxidation assay

Lipid peroxidation was estimated by quantifying malondialdehyde (MDA) using the thiobarbituric acid (TBA) method. Frozen tissue (0.25 g) was homogenized in 0.1% (w/v) TCA and centrifuged as described above.

An aliquot of the supernatant was reacted with TBA solution (0.5% in 20% TCA) and incubated at high temperature for 30 min. The reaction was terminated by rapid cooling on ice. Absorbance was measured at 532 nm, and non-specific turbidity was corrected by subtracting readings at 600 nm. MDA concentration was calculated using an extinction coefficient of 155 mM^−1^ cm^−1^.

### 2.5. Determination of ascorbate pool components

Total ascorbate (AsA + DHA) and reduced ascorbate (AsA) contents were determined using a modified microplate protocol. Frozen samples (0.5 g) were extracted in cold 6% (w/v) TCA and centrifuged at 16,000 × g for 15 min at 4 °C.

For total ascorbate quantification, extracts were treated with dithiothreitol (DTT) to reduce dehydroascorbate (DHA), followed by neutralization with N-ethylmaleimide (NEM). Subsequently, a colorimetric reaction mixture containing orthophosphoric acid, bipyridyl, and ferric chloride was added. After incubation at 42 °C, absorbance was recorded at 525 nm.

AsA content was measured using the same procedure without DTT and NEM. DHA concentration was calculated as the difference between total ascorbate and reduced ascorbate. Standard curves were established using L-ascorbic acid.

### 2.6. Gene expression analysis

Total RNA was extracted from frozen fruit tissues using Tri Reagent according to the manufacturer’s protocol. RNA quantity and purity were assessed spectrophotometrically (A260/A280), and integrity was verified by agarose gel electrophoresis.

First-strand cDNA synthesis was performed from 4 µg of total RNA using an oligo(dT) primer system following thermal denaturation and reverse transcription steps. Quantitative real-time PCR (qRT-PCR) was conducted using SYBR Green chemistry on a Realplex system (Eppendorf).

Gene expression levels were normalized against the reference gene SlActin, and relative transcript abundance was calculated accordingly. Gene-specific primers targeting antioxidant-related genes were designed based on sequences retrieved from the GenBank database using DNAMAN software. PCR products were purified, sequenced, and deposited in GenBank under the corresponding accession numbers.

### 2.7. Antioxidant enzyme activities

#### 2.7.1. PROTEIN extraction

Proteins were extracted from frozen fruit tissues homogenized in MES/KOH buffer (50 mM, pH 6.0) containing KCl, CaCl_2_, and ascorbate. After centrifugation at 16,000 × g for 15 min at 4 °C, the supernatants were collected for enzymatic assays. Protein concentration was determined using the Bradford method.

### 2.7.2. Enzyme assays

Enzyme activities were measured spectrophotometrically in microplate-based kinetic assays at 25 °C. Activities of APX, DHAR, MDHAR, and GR were determined according to established protocols.

Superoxide dismutase (SOD) activity was evaluated based on its ability to inhibit the photochemical reduction of nitro blue tetrazolium (NBT) under light exposure, with absorbance measured at 560 nm.

Catalase (CAT) activity was determined by monitoring the decomposition rate of hydrogen peroxide at 240 nm. Enzyme activities were calculated using appropriate extinction coefficients and expressed relative to protein content.

### 2.8. Statistical analysis

All experiments were conducted in triplicate. Data are presented as mean values ± standard error (SE). Statistical differences among treatments were evaluated by analysis of variance (ANOVA) using MINITAB software, with significance set at p < 0.05.

## 3. Results

### 3.1. Plant water status

Mercury exposure resulted in a pronounced reduction in predawn leaf water potential (LΨw), with the magnitude of decline being both concentration- and time-dependent (Table 2). In contrast, no significant alterations were observed in fruit water potential (FΨw), fruit osmotic potential (FΨO), or water content across fruits, leaves, and peduncles (FWC, LWC, PWC) (Table 2).

**Table 1.**
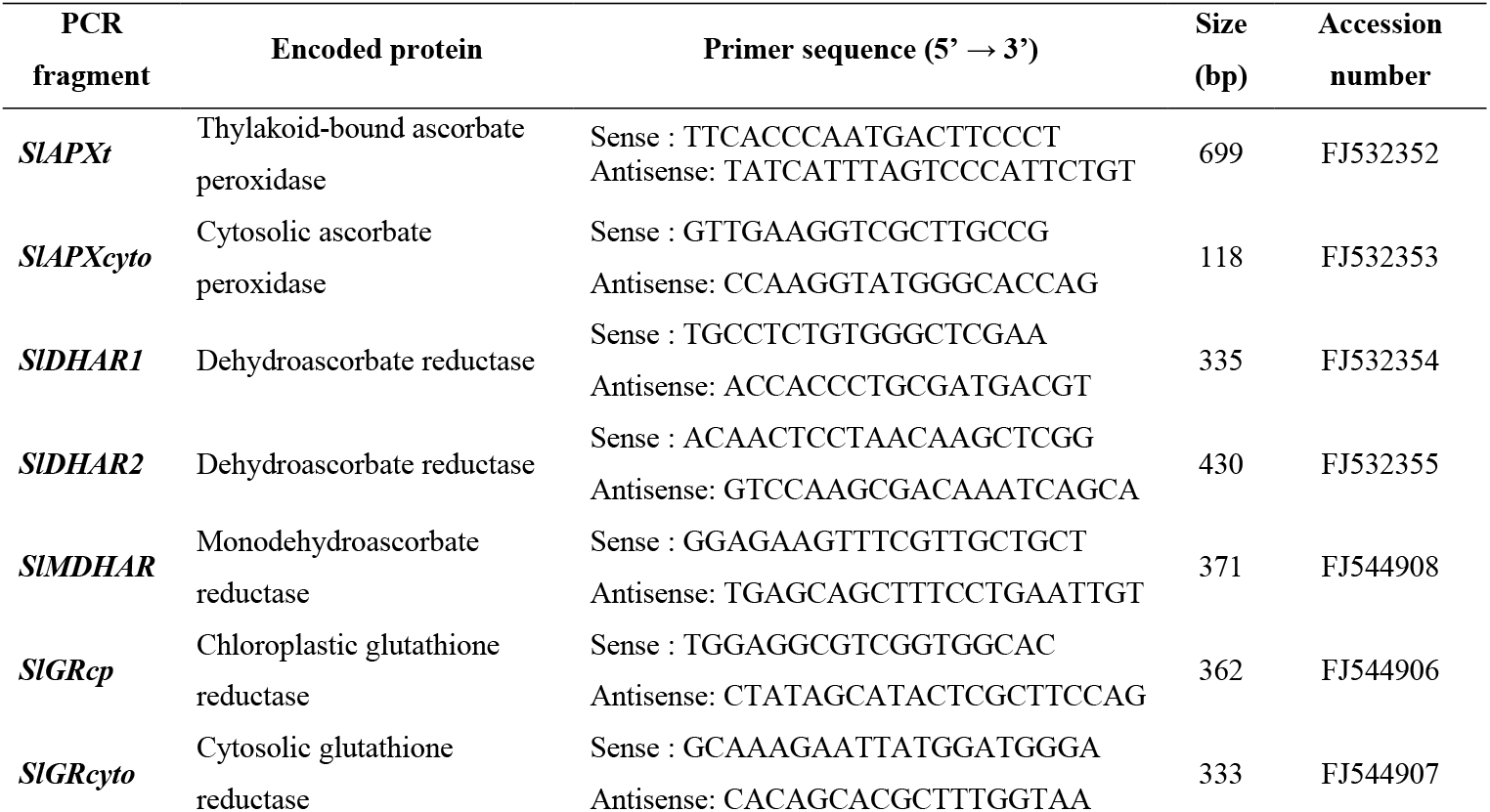
Sets of primers used to amplify gene-specific regions, corresponding size and accession number of the amplified product.

**Table 2.** Predawn leaf water potential (LΨ_w_; MPa), fruit water potential (FΨ_w_; MPa), fruit osmotic potential (FΨ_O_; MPa) and water content of fruits (FWC; %), leaves (LWC; %) and fruit peduncles (PWC; %) of control plants (C) and plants treated with 2, 5, 10, and 20 ppm of HgCl_2_ for 24 and 48 hours. Values are the mean (± S.E.) of five replicates, and different letters within lines indicate significant differences (P<0.05).

|  | 24 h |  |  |  |  | 48 h |  |  |  |  |
| --- | --- | --- | --- | --- | --- | --- | --- | --- | --- | --- |
|  | C | 2 ppm | 5 ppm | 10 ppm | 20 ppm | C | 2 ppm | 5 ppm | 10 ppm | 20 ppm |
| $L\Psi_w$ | $-0.34 \pm 0.02^a$ | $-0.48 \pm 0.02^b$ | $-0.63 \pm 0.03^c$ | $-1.15 \pm 0.07^d$ | $-1.37 \pm 0.06^e$ | $-0.35 \pm 0.02^a$ | $-0.45 \pm 0.02^b$ | $-0.65 \pm 0.03^c$ | $-1.35 \pm 0.03^e$ | $-1.60 \pm 0.04^f$ |
| $F\Psi_w$ | $-0.45 \pm 0.04^a$ | $-0.38 \pm 0.09^a$ | $-0.48 \pm 0.10^a$ | $-0.41 \pm 0.02^a$ | $-0.42 \pm 0.09^a$ | $-0.44 \pm 0.03^a$ | $-0.42 \pm 0.05^a$ | $-0.45 \pm 0.03^a$ | $-0.43 \pm 0.08^a$ | $-0.46 \pm 0.05^a$ |
| $F\Psi_o$ | $-0.72 \pm 0.01^a$ | $-0.68 \pm 0.03^a$ | $-0.77 \pm 0.03^a$ | $-0.83 \pm 0.01^a$ | $-0.82 \pm 0.02^a$ | $-0.73 \pm 0.02^a$ | $-0.76 \pm 0.01^a$ | $-0.78 \pm 0.02^a$ | $-0.78 \pm 0.03^a$ | $-0.75 \pm 0.01^a$ |
| FWC | $94 \pm 0.5^a$ | $94 \pm 0.4^a$ | $94 \pm 0.5^a$ | $95 \pm 0.4^a$ | $94 \pm 0.2^a$ | $94 \pm 0.3^a$ | $95 \pm 0.4^a$ | $94 \pm 0.4^a$ | $94 \pm 0.6^a$ | $95 \pm 0.3^a$ |
| LWC | $91 \pm 0.4^a$ | $91 \pm 1.8^a$ | $92 \pm 0.6^a$ | $91 \pm 0.3^a$ | $90 \pm 0.3^a$ | $91 \pm 0.4^a$ | $90 \pm 0.2^a$ | $90 \pm 0.4^a$ | $90 \pm 0.8^a$ | $89 \pm 1.5^a$ |
| PWC | $86 \pm 5.0^a$ | $89 \pm 4.5^a$ | $86 \pm 0.8^a$ | $82 \pm 5.6^a$ | $84 \pm 0.4^a$ | $86 \pm 4.2^a$ | $83 \pm 2.4^a$ | $83 \pm 0.2^a$ | $83 \pm 0.6^a$ | $82 \pm 0.8^a$ |

Water transport dynamics were, however, affected by mercury, as evidenced by a progressive decline in water flux up to 10 ppm HgCl_2_, followed by a complete cessation at 20 ppm, indicating a threshold beyond which hydraulic conductivity is severely impaired.

### 3.2. Oxidative markers: H_2_O_2_ and lipid peroxidation

Mercury treatments triggered a marked accumulation of H_2_O_2_ in leaves, particularly at moderate to high concentrations during early exposure (24 h), while prolonged exposure (48 h) led to a decline at the highest concentration tested (Table 3).

**Table 3.** H_2_O_2_ (nmol. g^-1^ FW) and malonyldialdehyde (MDA; nmol. g^-1^ FW) contents in leaves and fruits of control plants (C) and plants treated with 2, 5, 10, and 20 ppm of HgCl_2_ for 24 and 48 hours. Values are the mean (± S.E.) of five replicates, and different letters within columns indicate significant differences (P<0.05).

|  | Leaves |  |  |  | Fruits |  |  |  |
| --- | --- | --- | --- | --- | --- | --- | --- | --- |
|  | 24h |  | 48h |  | 24h |  | 48h |  |
| | $H_2O_2$ | MDA | $H_2O_2$ | MDA | $H_2O_2$ | MDA | $H_2O_2$ | MDA |
| C | $849.36 \pm 17.64^a$ | $34.04 \pm 0.19^a$ | $850.31 \pm 16.48^a$ | $34.25 \pm 0.27^a$ | $56.77 \pm 3.24^a$ | $34.74 \pm 5.24^a$ | $58.82 \pm 3.29^a$ | $33.59 \pm 4.37^a$ |
| 2 ppm | $804.19 \pm 34.19^a$ | $33.19 \pm 0.95^a$ | $856.76 \pm 37.30^a$ | $37.14 \pm 0.71^b$ | $51.47 \pm 4.41^a$ | $34.37 \pm 6.83^a$ | $55.29 \pm 49.71^a$ | $33.60 \pm 5.97^a$ |
| 5 ppm | $1081.70 \pm 107.62^b$ | $33.36 \pm 0.53^a$ | $1063.76 \pm 29.65^c$ | $38.92 \pm 0.66^b$ | $54.41 \pm 7.06^a$ | $35.13 \pm 6.62^a$ | $55.30 \pm 6.47^a$ | $35.08 \pm 5.41^a$ |
| 10 ppm | $1013.49 \pm 87.27^b$ | $32.16 \pm 1.42^a$ | $1212.60 \pm 22.26^d$ | $43.51 \pm 0.68^c$ | $57.35 \pm 2.47^a$ | $37.04 \pm 3.21^a$ | $59.71 \pm 5.88^a$ | $39.39 \pm 3.03^a$ |
| 20 ppm | $1211.88 \pm 93.64^b$ | $48.67 \pm 0.65^b$ | $476.93 \pm 21.36^e$ | $44.26 \pm 1.57^c$ | $58.53 \pm 2.35^a$ | $35.09 \pm 5.66^a$ | $51.18 \pm 4.94^a$ | $34.65 \pm 6.90^a$ |

Lipid peroxidation, assessed via MDA levels, showed a clear induction under mercury stress, especially after prolonged exposure (Table 3). Notably, these oxidative responses were restricted to leaf tissues, as fruit tissues did not exhibit significant changes in either H_2_O_2_ or MDA levels (Table 3), suggesting tissue-specific sensitivity to mercury-induced oxidative stress.

### 3.3. Non-enzymatic antioxidant responses

The ascorbate pool exhibited differential modulation depending on tissue type and exposure duration. In leaves, reduced ascorbate (AsA) levels increased under most mercury treatments, while dehydroascorbate (DHA) generally declined during early exposure but displayed an opposite trend after prolonged treatment. Consequently, the ascorbate redox state was enhanced under short-term stress conditions (Table 4).

**Table 4.** Total ascorbate, Ascorbate (AsA) and dehydroascorbate (DHA) concentrations (nmol. g^-1^ FW) and ascorbate redox state (AsA/Total) in leaves and fruit of control plants (C) and plants treated with 2, 5, 10, and 20 ppm of HgCl_2_ for 24 and 48 hours. Values are the mean (± S.E.) of five replicates, and different letters within columns indicate significant differences (P<0.05).

|  | Leaves |  |  |  |  |  | Fruits |  |  |  |  |  |
| --- | --- | --- | --- | --- | --- | --- | --- | --- | --- | --- | --- | --- |
|  | 24h |  |  | 48h |  |  | 24h |  |  | 48h |  |  |
|  | AsA | DHA | Redox state | AsA | DHA | Redox state | AsA | DHA | Redox state | AsA | DHA | Redox state |
| C | 27.50 ± 1.65<sup>a</sup> | 24.41 ± 1.13<sup>a</sup> | 0.53 ± 0.03<sup>a</sup> | 27.50 ± 1.42<sup>a</sup> | 24.38 ± 1.08<sup>a</sup> | 0.52 ± 0.03<sup>a</sup> | 21.07 ± 1.33<sup>a</sup> | 2.63 ± 0.85<sup>a</sup> | 0.89 ± 0.03<sup>a</sup> | 21.13 ± 1.29<sup>a</sup> | 2.65 ± 0.82<sup>a</sup> | 0.89 ± 0.03<sup>a</sup> |
| 2 ppm | 32.49 ± 2.15<sup>a</sup> | 20.11 ± 1.75<sup>b</sup> | 0.62 ± 0.02<sup>b</sup> | 55.19 ± 2.65<sup>b</sup> | 20.98 ± 1.15<sup>b</sup> | 0.73 ± 0.04<sup>b</sup> | 18.21 ± 2.81<sup>a</sup> | 2.07 ± 0.50<sup>a</sup> | 0.90 ± 0.02<sup>a</sup> | 14.26 ± 1.07<sup>b</sup> | 5.83 ± 0.69<sup>b</sup> | 0.71 ± 0.02<sup>b</sup> |
| 5 ppm | 41.60 ± 1.51<sup>b</sup> | 13.96 ± 1.22<sup>c</sup> | 0.75 ± 0.03<sup>c</sup> | 43.73 ± 1.44<sup>c</sup> | 34.68 ± 1.24<sup>c</sup> | 0.56 ± 0.04<sup>a</sup> | 20.40 ± 1.76<sup>a</sup> | 2.61 ± 0.39<sup>a</sup> | 0.89 ± 0.03<sup>a</sup> | 16.36 ± 1.28<sup>b</sup> | 6.87 ± 0.47<sup>b</sup> | 0.70 ± 0.03<sup>b</sup> |
| 10 ppm | 43.61 ± 2.17<sup>b</sup> | 11.29 ± 1.34<sup>c</sup> | 0.79 ± 0.04<sup>c</sup> | 43.53 ± 1.43<sup>c</sup> | 35.30 ± 1.42<sup>c</sup> | 0.55 ± 0.02<sup>a</sup> | 20.45 ± 3.62<sup>a</sup> | 2.37 ± 0.42<sup>a</sup> | 0.93 ± 0.03<sup>a</sup> | 14.44 ± 0.20<sup>b</sup> | 6.63 ± 0.40<sup>b</sup> | 0.69 ± 0.02<sup>b</sup> |
| 20 ppm | 41.79 ± 2.05<sup>b</sup> | 17.28 ± 1.23<sup>b</sup> | 0.71± 0.03<sup>c</sup> | 48.56 ± 1.73<sup>d</sup> | 34.46 ± 0.90<sup>c</sup> | 0.59 ± 0.03<sup>a</sup> | 20.06 ± 1.81<sup>a</sup> | 2.59 ± 0.99<sup>a</sup> | 0.89 ± 0.02<sup>a</sup> | 13.59 ± 1.37<sup>b</sup> | 6.78 ± 0.33<sup>b</sup> | 0.67 ± 0.02<sup>b</sup> |

In fruits, however, AsA levels remained stable during early exposure but declined significantly after 48 h, accompanied by an increase in DHA and a marked reduction in the ascorbate redox state (Table 4). These findings indicate a shift toward a more oxidized intracellular environment in fruits under prolonged mercury exposure.

### 3.4. Expression of antioxidant-related genes

The transcriptional profiles of genes associated with the ascorbate–glutathione cycle revealed complex and concentration-dependent regulation patterns (Table 5). While moderate mercury levels generally stimulated the expression of genes such as SlAPXt, SlDHAR1, SlDHAR2, and SlMDHAR, higher concentrations and longer exposure periods led to transcriptional repression.

**Table 5.**
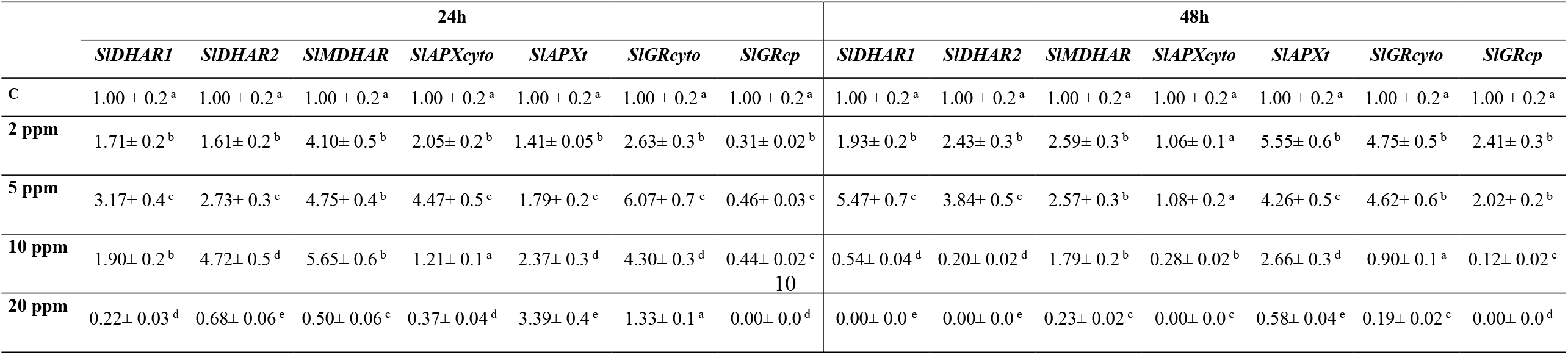
Changes in the levels of mRNA encoding antioxidant enzymes in fruits of control plants (C) and plants treated with 2, 5, 10, and 20 ppm of HgCl_2_ for 24 and 48 hours. Data were expressed as relative values with respect to the value found in the control fruits. Each value is the mean of nine replicates, and different letters within columns indicate significant differences (P<0.05).

Distinct isoform-specific responses were also observed. For instance, cytosolic and chloroplastic glutathione reductase genes (SlGRcyto and SlGRcp) displayed divergent expression patterns (Table 5), reflecting differential compartmental regulation of redox homeostasis.

### 3.5. Antioxidant enzyme activities

Mercury exposure induced a substantial activation of antioxidant enzymes in fruits. SOD activity increased consistently under all treatments, with a peak response at intermediate concentrations (Figure 1).

**Figure 1.**
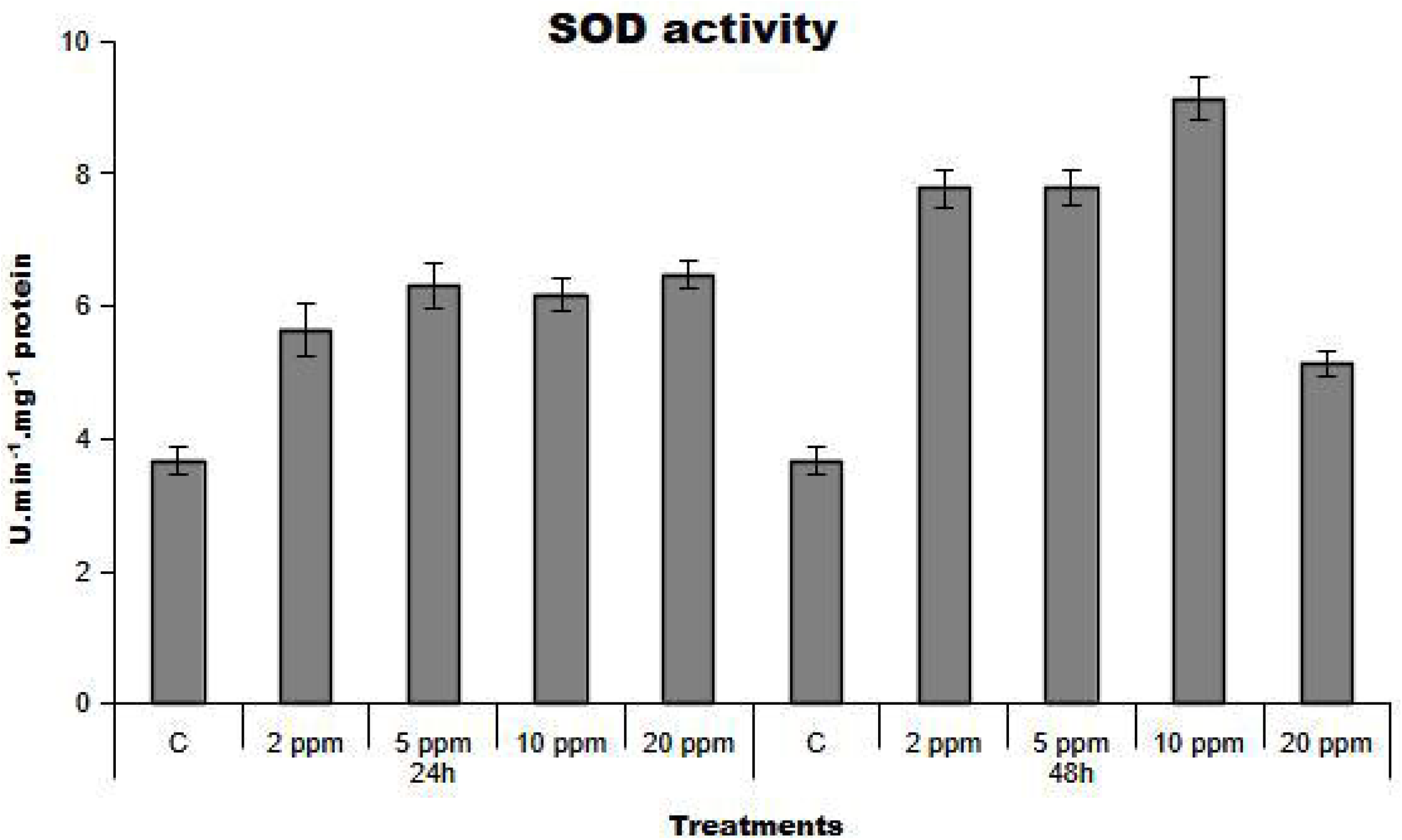
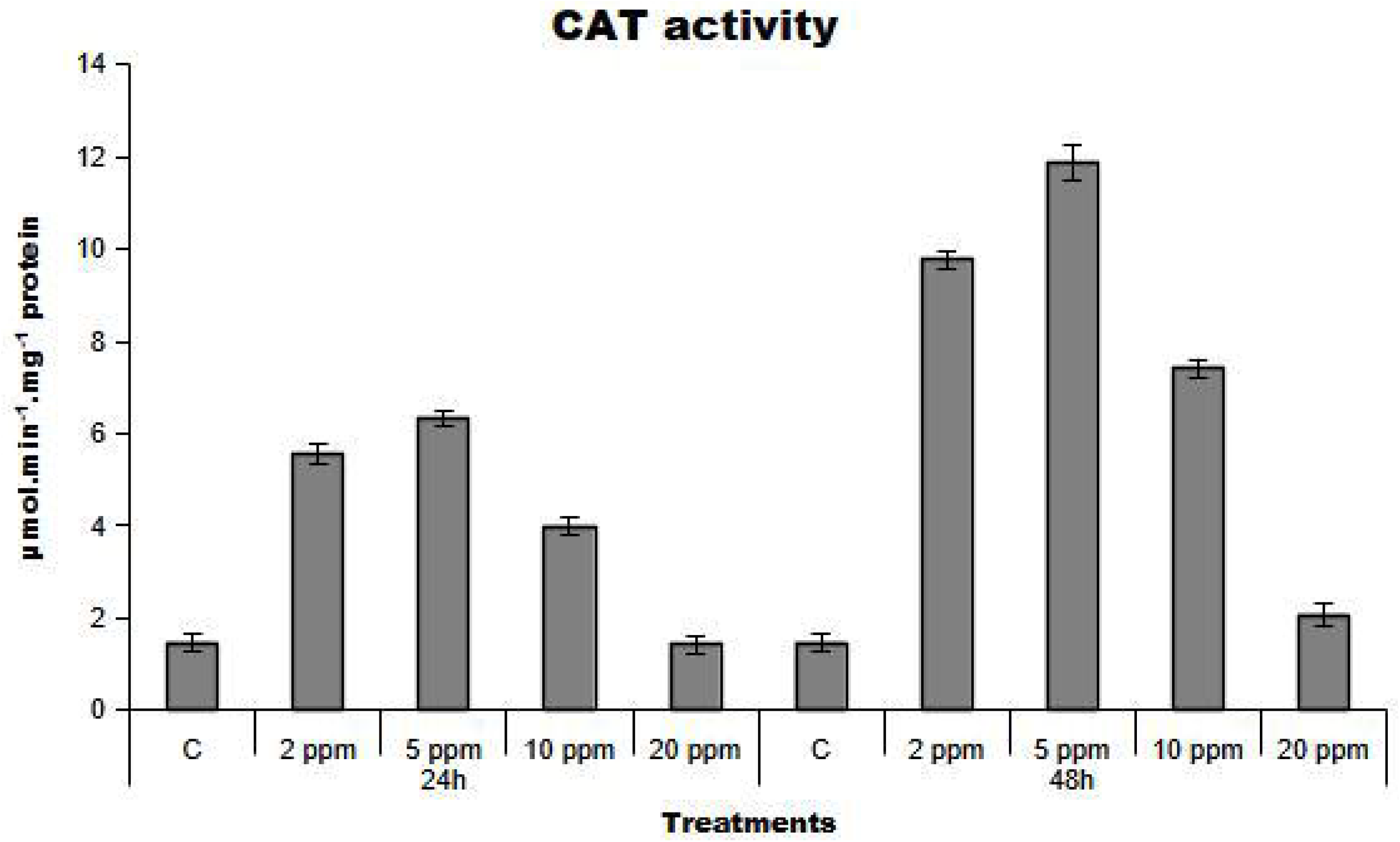
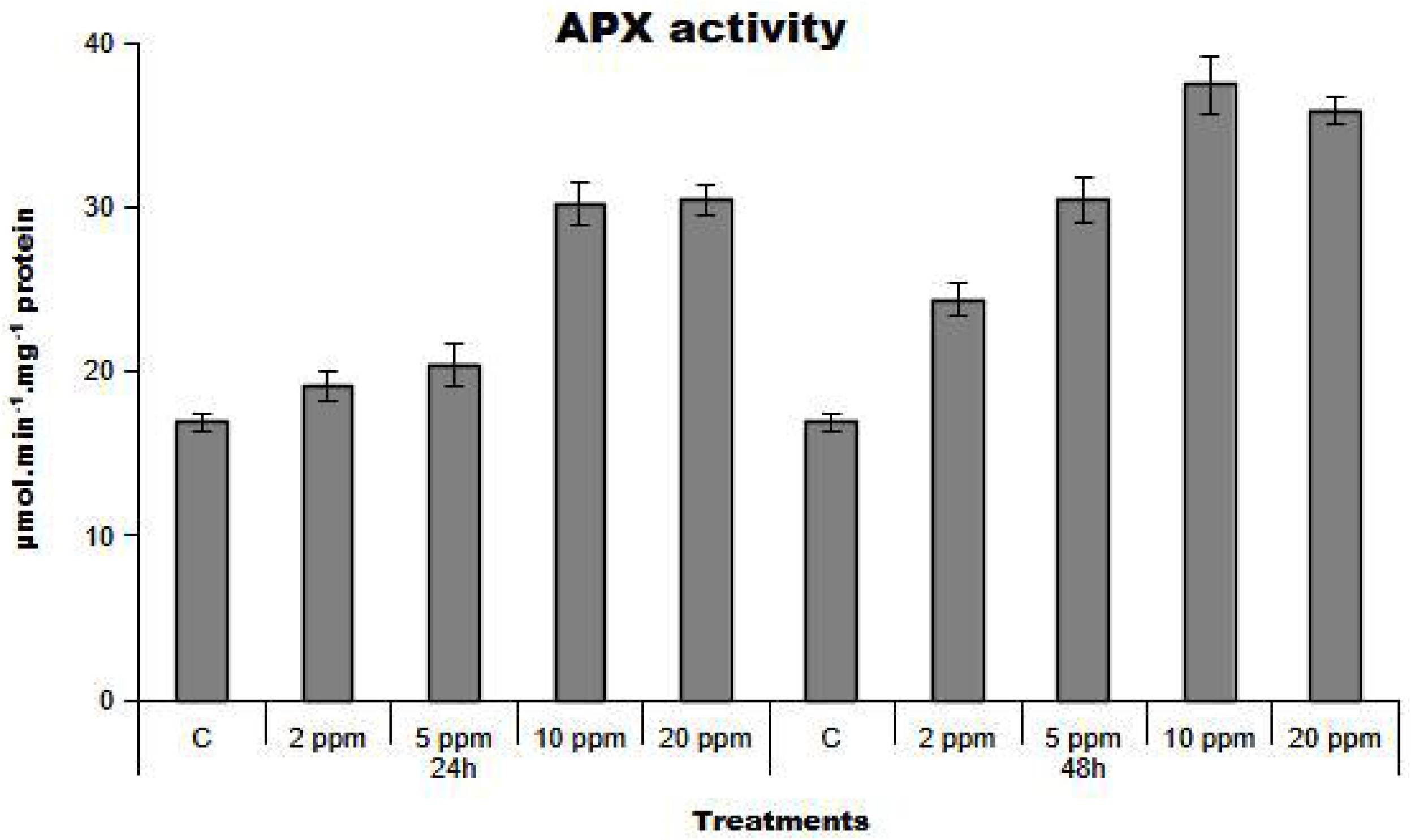
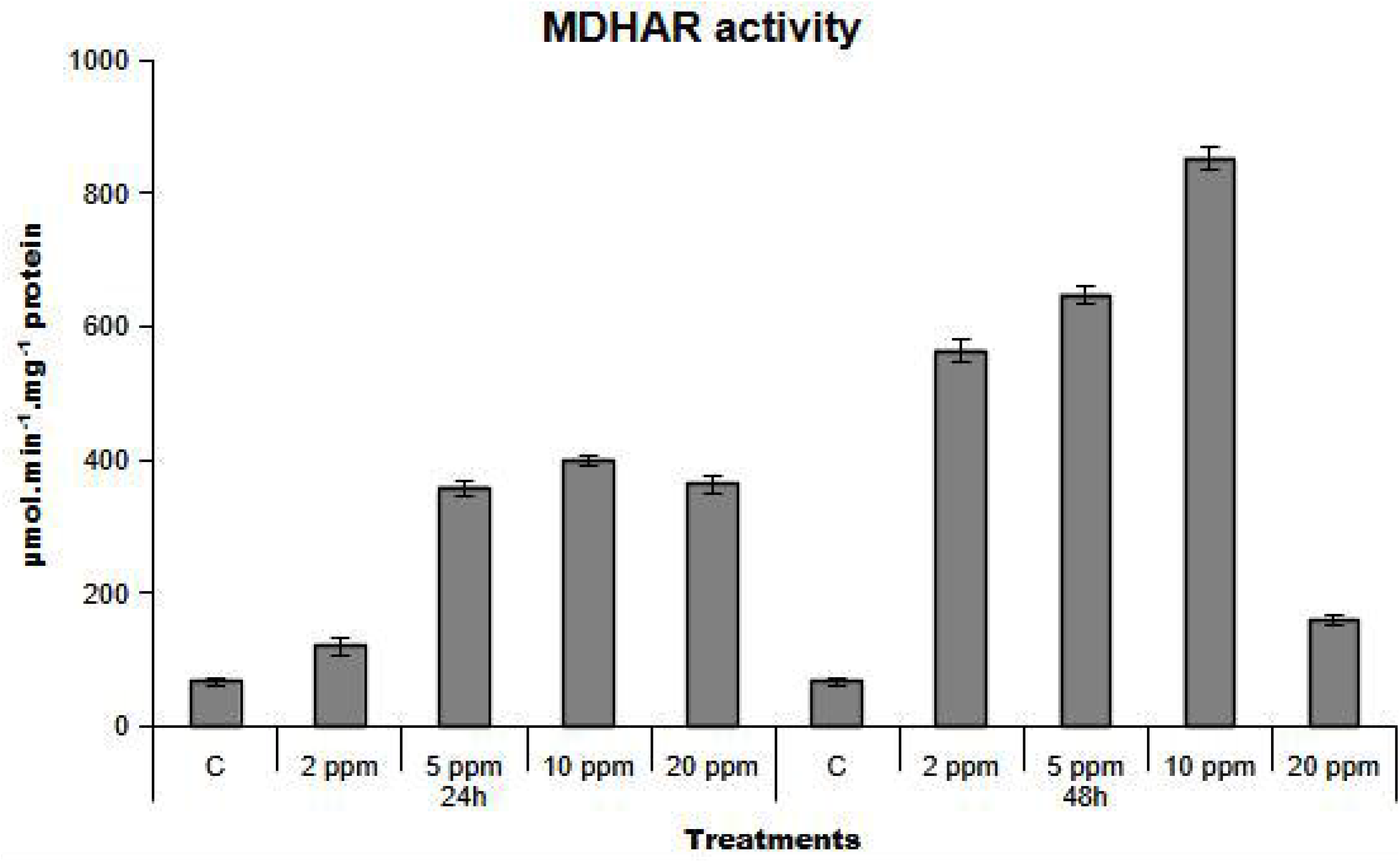
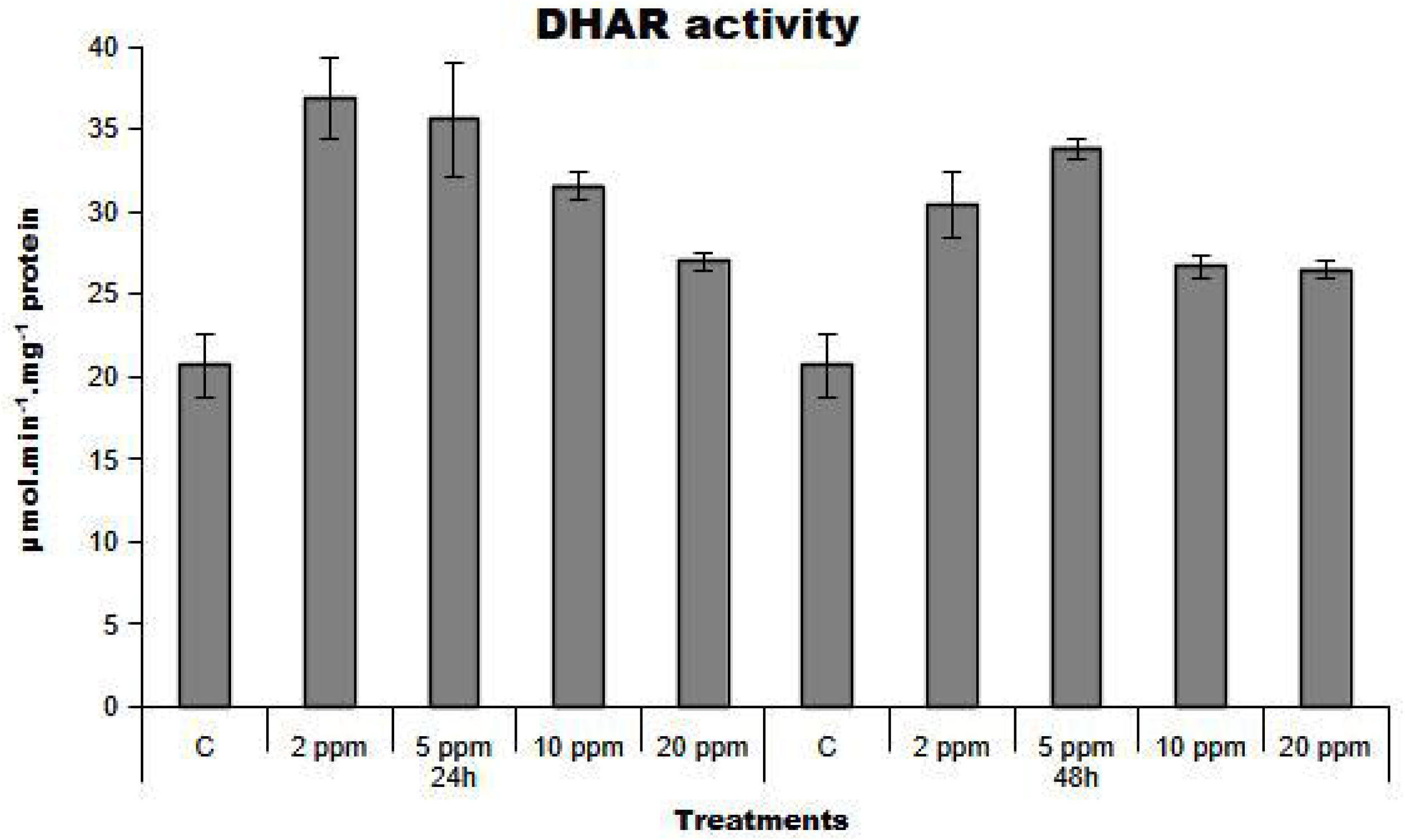
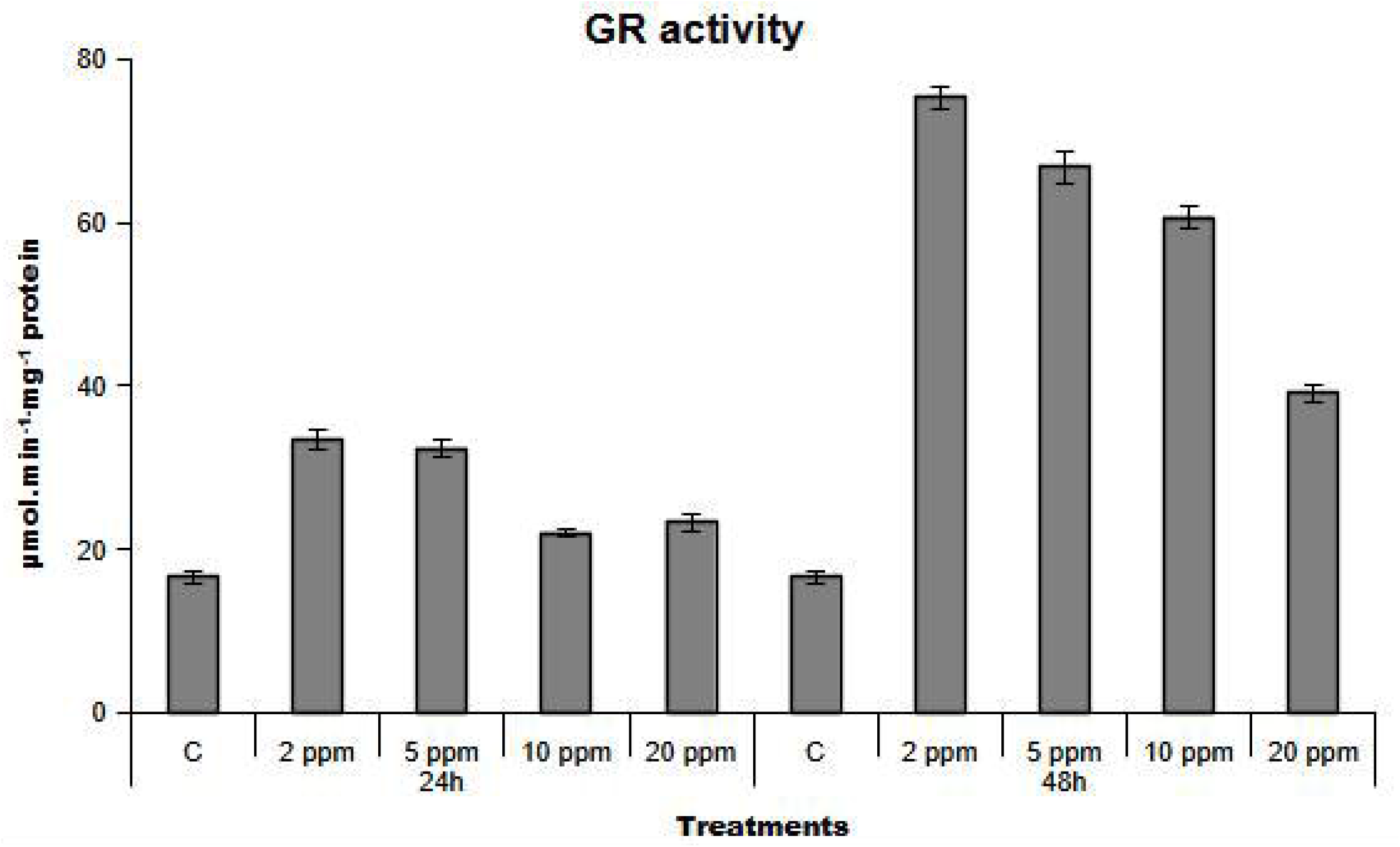
Activity of superoxide dismutase (SOD), catalase (CAT), ascorbate peroxidase (APX) monodehydroascobate reductase (MDHAR), dehydroascobate reductase (DHAR) and glutathione reductase (GR) in fruits of control plants (C) and plants treated by 2, 5, 10, and 20 ppm of HgCl_2_ for 24 and 48 hours.

CAT and APX activities were also strongly stimulated, indicating enhanced detoxification capacity for hydrogen peroxide (Figure 1). Enzymes involved in ascorbate recycling (MDHAR, DHAR, and GR) showed significant activation, although their responses varied with mercury concentration (Figure 1).

Notably, maximal induction was often observed at moderate concentrations, whereas higher concentrations resulted in partial inhibition, suggesting enzyme sensitivity to excessive mercury levels.

## 4. Discussion

The decline in leaf water potential observed under mercury exposure (Table 2) likely reflects impaired membrane transport processes, particularly those mediated by aquaporins, which are known targets of mercury inhibition. Despite this disruption at the leaf level, fruit water relations remained relatively stable (Table 2), indicating a degree of hydraulic buffering or compartmentalization.

The accumulation of reactive oxygen species in leaves, coupled with increased lipid peroxidation (Table 3), confirms that mercury induces oxidative stress primarily in vegetative tissues. The absence of similar changes in fruits (Table 3) suggests the existence of more efficient antioxidant systems in reproductive tissues, a hypothesis supported by the enhanced enzymatic activities observed (Figure 1).

The upregulation of SOD, CAT, and APX activities (Figure 1) indicates an increased flux through ROS detoxification pathways. However, the decline in enzyme efficiency at higher mercury concentrations suggests potential inhibitory effects of mercury on protein structure and function.

The ascorbate–glutathione cycle appears to play a central role, as reflected by changes in enzyme activities (Figure 1), gene expression patterns (Table 5), and ascorbate redox state (Table 4). Nevertheless, discrepancies between transcript levels and enzyme activities point toward additional regulatory mechanisms beyond transcriptional control.

In fruits, the reduction in ascorbate redox state under prolonged exposure (Table 4) reflects a shift toward oxidative conditions despite the activation of enzymatic defenses, indicating potential limitations in sustained stress tolerance.

## 5. Conclusions

Mercury exposure induces oxidative stress in tomato plants, as evidenced by increased ROS accumulation and lipid peroxidation in leaves (Table 3), alongside significant modulation of antioxidant systems (Figure 1; Table 4).

Although fruit tissues initially maintain redox stability, likely through efficient antioxidant responses, prolonged exposure results in measurable disturbances in redox balance (Table 4). These findings emphasize the importance of antioxidant mechanisms in mercury tolerance and highlight differential responses between plant organs.

## Acknowledgements

The authors gratefully acknowledges the financial support provided by the Scholar Rescue Fund (SRF) and the PAUSE Program (Programme d’aide à l’accueil en urgence des scientifiques et des artistes en exil), which made the publication of this article possible.

## Author contributions

Ramzi MURSHED : Conceptualization, Investigation, Methodology, Writing – original draft. Sanders JUNGLEE : Methodology. Huguette SALLANON : Supervision, Validation, Writing – review and editing. Laurent URBAN : Validation, Writing – review and editing. Félicie LAURI : Methodology, Project administration, Writing – review and editing.

